# Identification and characterization of a phase-variable element that regulates the autotransporter UpaE in uropathogenic *Escherichia coli*

**DOI:** 10.1101/356865

**Authors:** E.J. Battaglioli, K.G.K Goh, T. S. Atruksang, K. Schwartz, M. A. Schembri, R.A. Welch

## Abstract

Uropathogenic *Escherichia coli* (UPEC) are the most common etiological agent of uncomplicated urinary tract infection (UTI). An important mechanism of gene regulation in UPEC is phase variation that involves inversion of a promoter-containing DNA element via enzymatic activity of tyrosine recombinases, resulting in biphasic, ON or OFF expression of target genes. The UPEC reference strain CFT073 has five tyrosine site-specific recombinases that function at two previously characterized promoter inversion systems, *fimS* and *hyxS*. Three of the five recombinases are located proximally to their cognate target elements, which is typical of promoter inversion systems. The genes for the other two recombinases, IpuA and IpuB are located distal from these sites. Here, we identified and characterized a third phase variable invertible element in CFT073, *ipuS* located proximal to *ipuA* and *ipuB*. The inversion of *ipuS* is catalyzed by four of the five CFT073 recombinases. Orientation of the element drives transcription of a two-gene operon containing *ipuR*, a predicted LuxR-type regulator, and *upaE*, a predicted autotransporter. We show that the predicted autotransporter UpaE is surface-located and facilitates biofilm formation as well as adhesion to extracellular matrix proteins in a K-12 recombinant background. Consistent with this phenotype, the *ipuS* ON condition in CFT073 results in defective swimming motility, increased adherence to human kidney epithelial cells, and a positive competitive kidney colonization advantage in experimental mouse UTI infections. Overall, the identification of a third phase-switch in UPEC that is regulated by a shared set of recombinases describes a complex phase-variable virulence network in UPEC.

**Importance:** Uropathogenic *Escherichia coli* (UPEC) is the most common cause of urinary tract infection (UTI). ON versus OFF phase-switching by inversion of small DNA elements at two chromosome sites in UPEC regulates the expression of important virulence factors, including the type 1 fimbriae adhesion organelle. In this report, we describe a third invertible element, *ipuS*, in the UPEC reference strain CFT073. The inversion of *ipuS* controls the phase variable expression of *upaE*, an autotransporter gene that encodes a surface protein involved in adherence to extracellular matrix proteins and colonization of the kidneys in a murine model of UTI.

## Introduction

Urinary tract infections (UTIs) are one of the most common infections diagnosed in clinics and hospitals. Nearly 50% of women will experience a UTI in their lifetime with treatment costs exceeding $3.5 billion annually in the United States (1, 2). The most common etiological agent of uncomplicated UTIs are uropathogenic *Escherichia coli* (UPEC), which account for ~80% of reported infections (3). The predicted reservoir of UPEC is the colon and infection follows an ascending route, which is initiated via colonization of the urethra. Bacteria that gain access to the urinary tract face a variety of host defense mechanisms including shedding of uroepithelial cells, low iron levels, rapid recruitment of phagocytes, host-derived antimicrobial peptides, and the cleansing flow of urine (4–11). Additionally, recent characterization of a urinary tract-specific microbiome suggests there may also be microbial barriers to infection as is observed in the gut (12, 13). To establish and maintain an infection, UPEC possess specialized virulence factors to overcome these defense mechanisms. Well-described examples include adhesive fimbriae, multiple iron acquisition systems, a polysaccharide capsule, effective reactive nitrogen species detoxification systems, and toxins such as hemolysin (14–21).

Type 1 fimbriae are polytrichous hair-like projections expressed on the surface of UPEC cells (22). They mediate attachment and invasion of the bladder epithelium, are a key component of the “stick or swim” lifestyle choice, and are critical to the establishment and maintenance of infection in the murine model of UTI (14, 15, 23, 24). Type 1 fimbriae were also recently shown to facilitate adherence to colonic epithelial cells and persistence in the gut (25). The expression of Type 1 fimbriae is phase variable as a result of rearrangement of the invertible element or “switch” *fimS* that contains a promoter (26, 27). In *E. coli* K-12, inversion of *fimS* is catalyzed by the proximal encoded genes for the tyrosine site-specific recombinases FimB and FimE (28). In addition to the recombinases, multiple DNA binding proteins including IHF, LRP and H-NS interact with *fimS* to facilitate formation of the appropriate DNA conformation necessary for holiday junction formation and recombination (29–33). Associated changes in expression and activity of both the recombinases and the accessory DNA binding proteins alter switching kinetics and result in population wide changes of phase state (34–37). Additionally, crosstalk with genes from other adhesive fimbriae and specific environmental conditions including pH, osmolarity, temperature and metabolite availability are known to facilitate these population phase state biases (38–43). In total, these regulatory mechanisms are predicted to adapt a population phase to suit changing metabolic and environmental queues.

In addition to these methods of regulation, CFT073 has three additional tyrosine recombinases, FimX, IpuA and IpuB, which are conserved in many UPEC strains (23). FimX and IpuA are also capable of catalyzing inversion of *fimS* in CFT073 despite being located distal to *fimS* on the CFT073 chromosome (23). Typically, site-specific recombinases that mediate inversion of phase switches are encoded proximal to their sites of functionality, suggesting the existence of other switches local to the three UPEC specific recombinases (44). Recently, a second phase variable element, *hyxS*, was characterized proximal to *fimX* in CFT073 and another UPEC strain UTI89 (45). Inversion of *hyxS* regulates expression of *hyxR*, a LuxR-type regulator. Only FimX is capable of catalyzing inversion of this switch, and *hyxS* dependent expression of *hyxR* affects resistance to reactive nitrogen species and intracellular macrophage survival though the precise mechanisms underlying these affects remain to be characterized (45). Phase variable switching at *fimS* and *hyxS* has also been examined in UPEC strains from the globally disseminated multidrug resistant ST131 clone, which possess functional FimE and FimX recombinases (46).

Because there are known invertible DNA elements proximal to *fimB*, *fimE* and *fimX*, we sought to determine if a third phase switch existed proximal to *ipuA* and *ipuB*. Here we report the identification of a third phase variable switch in CFT073, *ipuS*, located adjacent to the *ipuA* and *ipuB* recombinase genes. The switch is bounded by a set of 7bp-inverted repeats and the recombination half sites share sequence similarity with the *fimS* and *hyxS* invertible elements. Transcriptional analysis identifies the presence of transcription start site in the element, and four of the five recombinases (FimB excluded) are able to independently catalyze *ipuS* inversion. Inversion of the element affects transcription of *ipuR*, a predicted LuxR-type regulator and *upaE*, a predicted autotransporter. Phenotypic characterization of UpaE reveals that it is exposed at the cell surface, and can facilitate biofilm formation as well as adhesion to human extracellular matrix proteins. Further analysis of *ipuS* inversion reveals that a locked-ON state results in a defect in swimming motility, increased adherence to kidney epithelial cells, and a 5-fold advantage in colonization of the kidneys at 72hpi. Overall, this work identifies an UPEC switch that controls the phase variable expression of UpaE, an autotransporter that may contribute to UPEC infection in the complex, diverse microenvironments of the urinary tract.

## Results

### Identification of a phase variable element *ipuS*

Previous studies identified inversion sites associated with the FimB, FimE and FimX tyrosine recombinases in CFT073 (23, 45). In most other characterized tyrosine recombinase-mediated phase variation systems, the recombinases are active on closely linked invertible DNA elements. Thus, we hypothesized that there would be an invertible element proximal to the *ipuA* and *ipuB* recombinase genes. Immediately 5’ of *ipuA* is a putative two-gene operon containing *ipuR* (encoding a predicted LuxR-type regulator) and *upaE* (encoding a predicted autotransporter protein). Further analysis of this DNA region reveals *ipuA* and *ipuR* are separated by a 317bp intergenic spacer with no predicted open reading frames (Fig. 1A). The size of the spacer is consistent with other promoter inversion systems, suggesting it may contain an invertible element. To test this, a chromosomal *ipuR-lacZ* transcriptional fusion was generated (strain WAM5009) to detect inversion events in this region. When a stationary phase LB-broth culture of WAM5009 was plated on MacConkey’s lactose medium, the reporter strain displayed a mixture of red and white colonies. The region containing the predicted invertible element was amplified by PCR from a red and white colony respectively, and sequenced by Sanger dideoxy chain termination. The DNA sequences from the two colony types revealed the presence of a 260bp invertible element, which we refer to as *ipuS* (Fig. 1B). The *ipuS* element is bounded by a pair of 7bp-inverted repeats with the distal-inverted repeat located within the annotated coding sequence of *ipuA* (Fig. 1B). In the OFF state, defined as lack of expression from the *ipuR-lacZ* transcriptional fusion, the full-length form of IpuA is produced. Upon inversion to the ON state, defined as expression of the *ipuR-lacZ* fusion, a truncation of the *ipuA* coding sequence occurs. The truncation removes 11 amino acids from the C-terminus of IpuA and generates a K-L substitution of the terminal amino acid (Fig. 1C). None of the four required RHRY active site residues for IpuA are altered by the truncation, suggesting the shortened form may retain catalytic activity (Fig. 1C).

**FIG 1.**
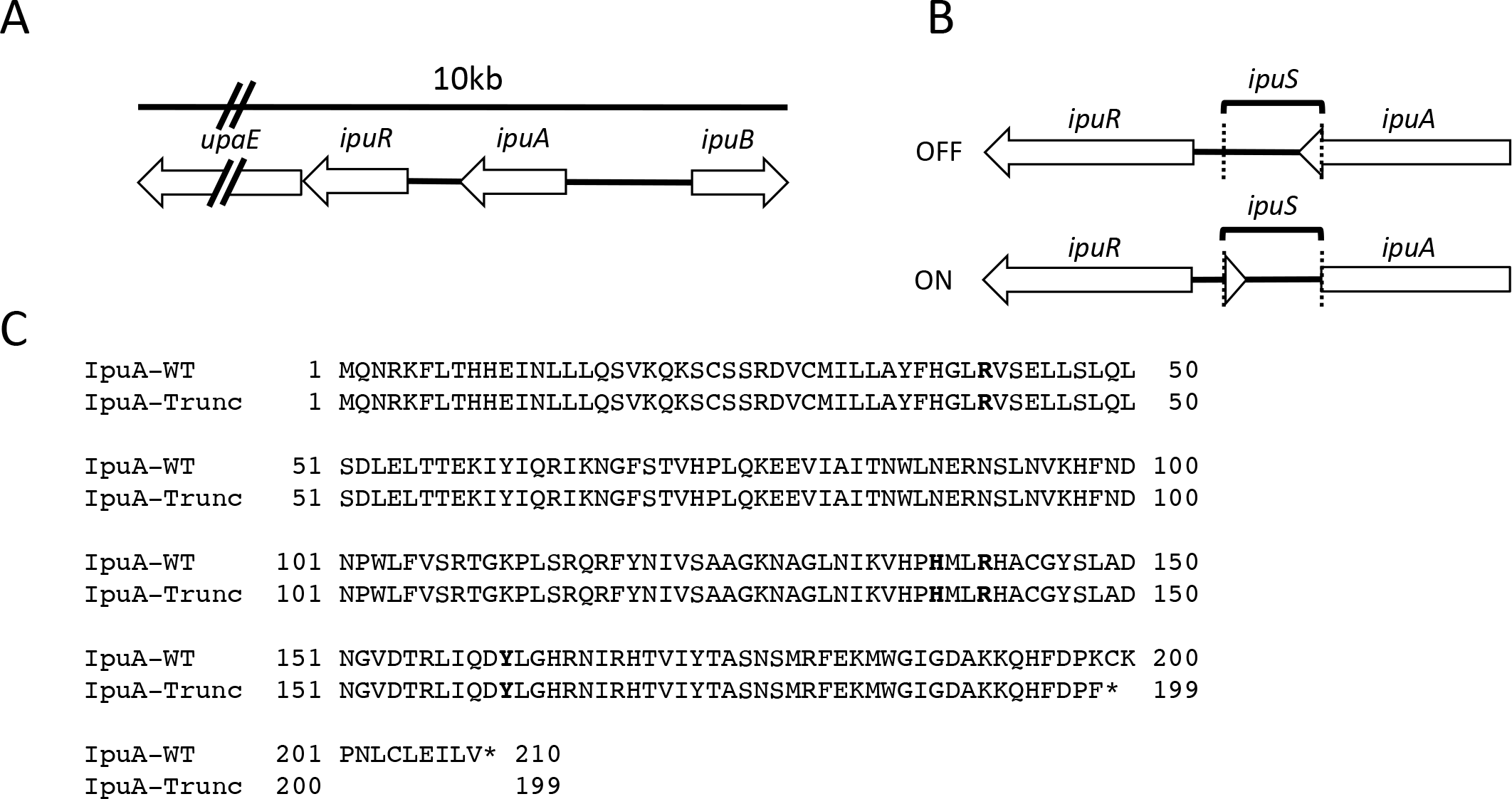
Identification of the *ipuS* invertible element. (A) Schematic representation of the genomic context of *ipuS.* (B) DNA rearrangement as a result of *ipuS* inversion. The ON/OFF state is defined by expression of a *lacZ-ipuR* transcriptional fusion in the pictured orientation. Location of the inverted repeats is indicated by the dotted lines. (C) Inversion to the ON state results in an 11 amino acid truncation of *ipuA* and a K-F substitution of the truncated forms terminal amino acid. Conserved RHRY active site residues are indicated in bold and unaffected by the truncation.

### *ipuS* half site analysis

The *ipuS* invertible element is defined by a pair of 7bp-inverted repeats (Fig. 2A). *ipuS* has the shortest inverted repeats of the three described elements in CFT073, with *fimS* and *hyxS* having 9 and 16bp repeats, respectively (27, 45). In addition to the core repeat sequence, up to 5bp surrounding the core can participate in base pairing and help facilitate inversion. The sequence of these residues is similar to the respective required regions of *fimS* (22) and the predicted hairpin structure generated during recombination illustrates these potential base-pairing interactions (Fig. 2B).

**FIG 2.**
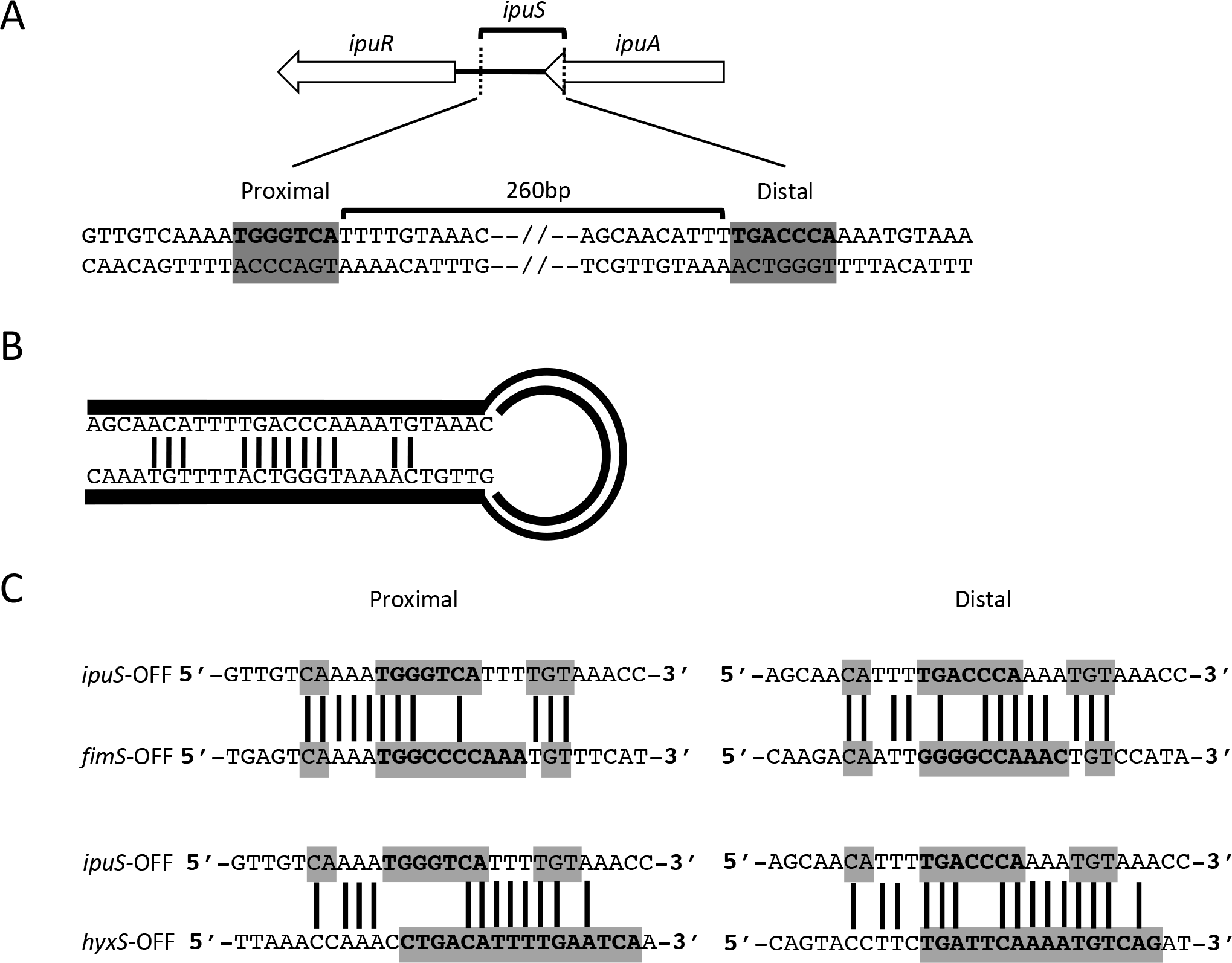
*ipuS* invertible element characterization. (A) The invertible element is 260bp and is defined by a pair of 7bp inverted repeats shown in bold with a grey background. (B) DNA hairpin created during recombination. Putative base pair interactions between the core inverted repeat and local flanking sequence are indicated by connecting lines. (C) Comparison of *ipuS* vs. *fimS* inverted repeat sequences (top panel) and *ipuS* vs. *hyxS* inverted repeat sequences (bottom panel). Proximal and distal half sites are shown on the left and right respectively. Inverted repeat sequence and surrounding sequences that can participate in base paring during recombination are boxed in grey and sequence identity is indicated by connecting bars. Nucleotides that are components of the core inverted repeats for each switch are shown in bold.

The inverted repeats and surrounding sequence of *ipuS* were compared to the same respective regions of *hyxS* and *fimS* to assess the potential of shared recombinase activity among the elements. For consistency, the OFF state of each element was used for the comparisons. The *ipuS* switch shares a high degree of sequence similarity with the other two switches, particularly *fimS* (Fig. 2C). In this figure, lines connecting bases indicate sequence identity; inverted repeat sequence is shown in bold and the grey boxes indicate potential base paring interactions. Previous reports comparing the half sites of *hyxS* and *fimS* show only limited similarity (45).

### Recombinase activity at *ipuS*

The activity of tyrosine recombinases at *fimS* and *hyxS* is known to be sequence specific, and the similarity between the *ipuS*, *fimS* and *hyxS* inverted repeats suggests that many if not all of the five recombinases would have activity at *ipuS*. To test this, the five recombinases were deleted by sequential Lambda-Red mutagenesis and ϕEB49 phage transduction to lock the orientation of all three switches. In the case of *ipuA*, a 404bp truncation from the 5’ end was generated to remove one of the required active site residues rendering the resulting protein nonfunctional, while preserving the *ipuS* distal inverted repeat and allowing for inversion via exogenous expression of the recombinases. Strains were created with all four possible combinations of *ipuS* and *fimS* phase states. *hyxS* was locked OFF in all strains examined (Table 1).

Each recombinase, including the full length (FL) and truncated (Trunc) forms of IpuA, was provided *in trans* on multicopy expression constructs in both the *ipuS* ON and OFF locked backgrounds. All the recombinase complementation plasmids were constructed in a pACYC177 background and constitutively expressed except for IpuA-FL and IpuA-Trunc. These were constructed in a pACYC184 background with the native *ipuA* promoter driving expression. In a previous publication, we showed that expression of *ipuA* under control of the kanamycin resistance gene promoter on pACYC177 causes cell morphology defects and expression from its native promoter rectifies this complication (23).

The ability of a single recombinase to switch the orientation from a starting ON or OFF state at *ipuS* was assayed by PCR amplification of the switch and asymmetric restriction digestion of the resulting product by PacI. With the exception of FimB, all the recombinases are independently capable of catalyzing inversion in both directions, including the truncated form of IpuA (Fig. 3). FimB showed no detectable catalytic activity under the conditions tested. However, the same pACYC177::*fimB* construct was capable of inverting *fimS*, demonstrating the lack of activity is not due to complications with recombinant plasmid expression (23). The inversion assay is not explicitly quantitative however inspection of the intensity of the bands in the digest suggests there may be differences in catalytic efficiency among the recombinases. FimE is less efficient than IpuA-FL, IpuA-Trunc, IpuB and FimX at inverting the *ipuS* switch in both directions under the conditions tested (Fig. 3). IpuB also displays a reduced capacity to catalyze ON to OFF inversion (Fig. 3).

**FIG 3.**
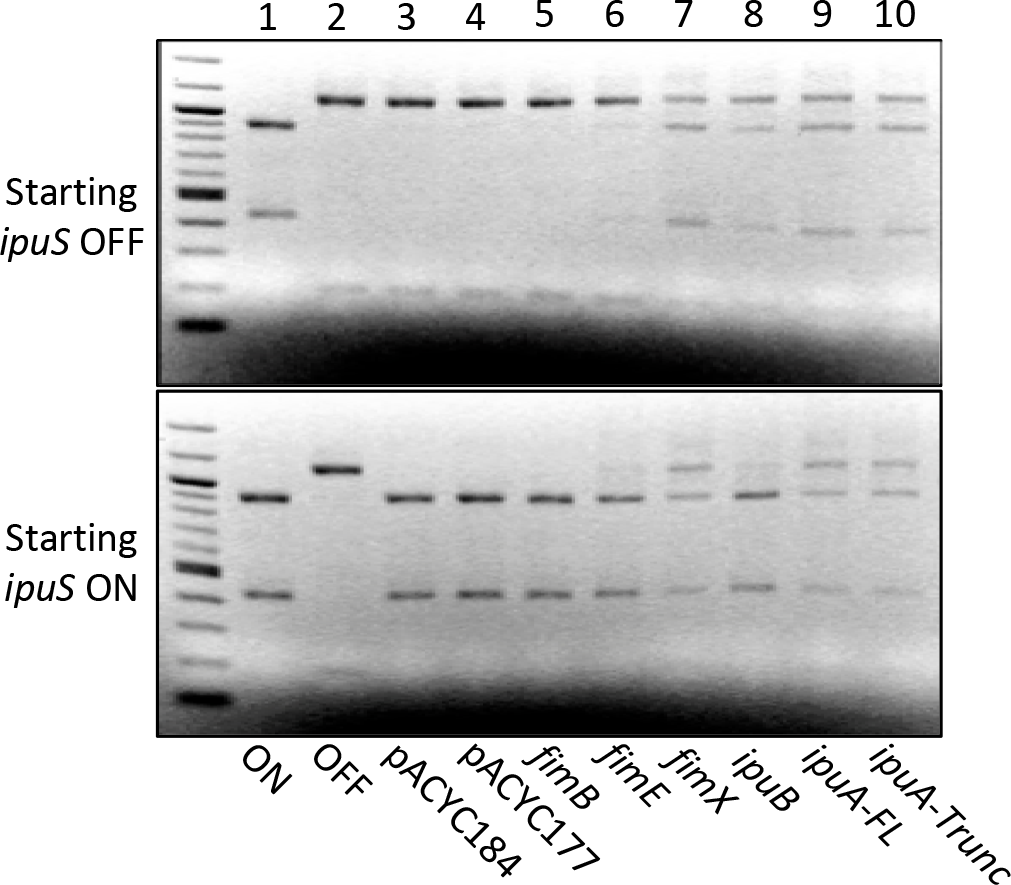
Assessment of each recombinases ability to catalyze inversion of *ipuS* in vitro. Ethidium bromide stained electrophoretic gels of Pad digested PCR products are shown. Lanes 1 and 2 contain digested PCR products from CFT073 *ipuS* phase lock ON (WAM5064) and OFF (WAM5065) strains generated by 5-way recombinase deletion. Lanes 3 and 4 contain digested PCR products from vector only controls (WAM5070, WAM5079, WAM5074 and WAM5083). Lanes 5-10 contain PCR products from the locked OFF strain WAM5065 (top panel) or locked ON strain WAM5064 (bottom panel) after transforming each with a recombinant plasmid containing the indicated recombinase. Both the full length and truncated forms of *ipuA* were tested for activity (lanes 9 and 10).

### Identification of a putative promoter in *ipuS*

We postulated that the invertible element could regulate transcription of *ipuR-upaE* by containing an additional promoter, or by blocking read through of an upstream *ipuA*-associated promoter. To test this, we subjected the *ipuS* region to 5’RACE using cDNA generated from *ipuS* phase locked ON and OFF strains. Only the locked-ON strain generated a product and subsequent sequencing revealed the location of a putative transcriptional start site in *ipuS* (Fig. 4A). Sequence analysis immediately upstream of the mapped transcriptional start site revealed a putative promoter with −35 and −10 sequences that each have 4 of 6 nucleotides matching the sigma-70 consensus sequence (Fig. 4B) (47).

**FIG 4.**
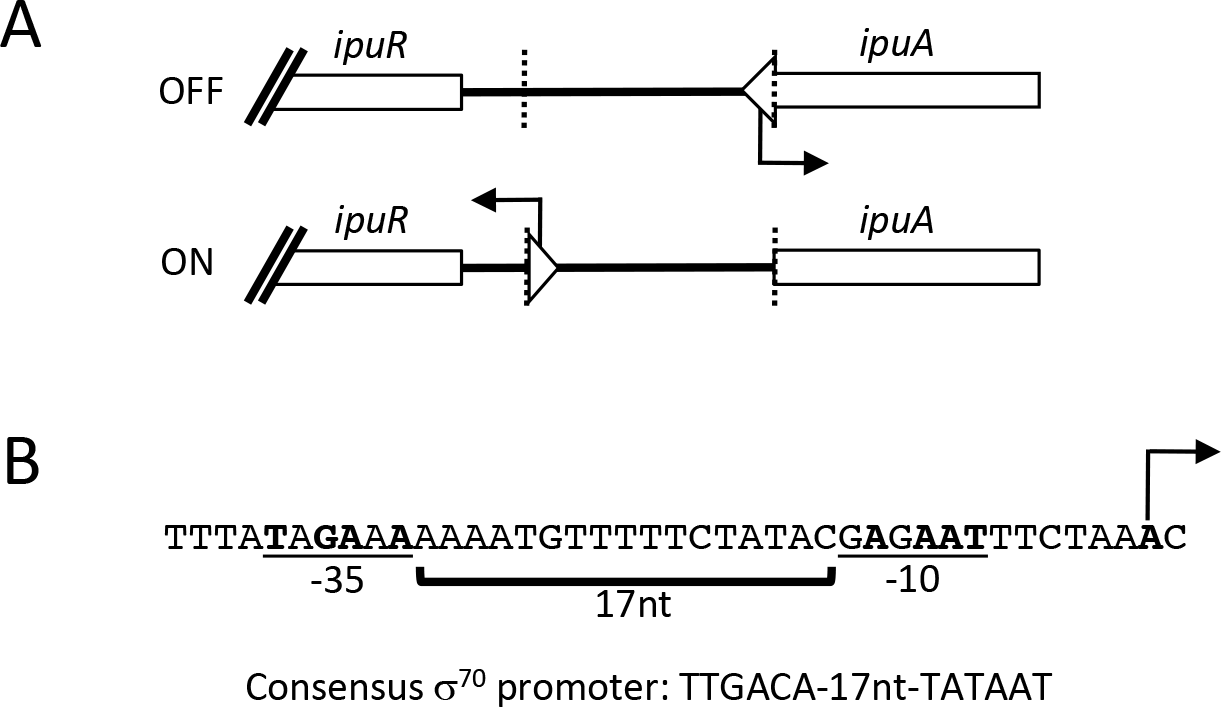
*ipuS* promoter mapping by 5’RACE. (A) The *ipuS* transcription start site was identified by 5’RACE and is located within the full length *ipuA* coding sequence. Inversion of element to the “ON” state orients the promoter toward *ipuR/upaE.* The inverted repeats are indicate by the dotted lines. (B) A predicted promoter was identified in proximity to the mapped transcription start site that most closely matches a s^70^ consensus promoter. Locations of the predicted −35 and −10 regions, spacer lengths and transcription start side are indicated. Bases matching consensus are shown in bold.

### UpaE is localized to the cell surface

We next sought to characterize phenotypic effects of the *ipuS* ON versus OFF phase state and began by assessing the functionality of the regulated genes *upaE* and *ipuR*. Initial genetic studies examining the role of *ipuR* did not reveal any clear phenotype, so we focused on characterization of the predicted autotransporter gene *upaE*. To assess the functionality of UpaE in isolation, we utilized a plasmid based overexpression system it the *E. coli* K-12 background strain MS427 (48). MS427 has a mutation in the Ag43-encoding *flu* gene rendering it unable to facilitate biofilm formation or self-aggregation, and has previously been used successfully to probe the function of other autotransporters (20, 48–53). Immunoblots of whole cell lysates generated from MS427 transformed with a UpaE expression plasmid using a polyclonal anti-serum raised to a UpaE-MBP fusion protein showed a band consistent with the 271kDa predicted molecular weight of UpaE (Fig. 5A). UpaE localization was then assessed using immunofluorescence microscopy which showed staining concentrated to the cell membrane suggesting it is membrane bound (Fig. 5B). Further assessment in the native CFT073 context yielded similar results. We probed for the expression of UpaE in whole cell lysates and culture supernatants of the phase lock ON and OFF CFT073 strains. UpaE was only detectable in phase lock ON cells (Fig. 5C). Additionally, extracellular UpaE species were not detected in concentrated 10ml TCA preparations from culture supernatants of either lock ON or OFF strains (Fig. 5C). This further corroborates the immunofluorescence data in the MS427 background and suggest that UpaE is membrane associated in the native CFT073 context (Fig. 5C).

**FIG 5.**
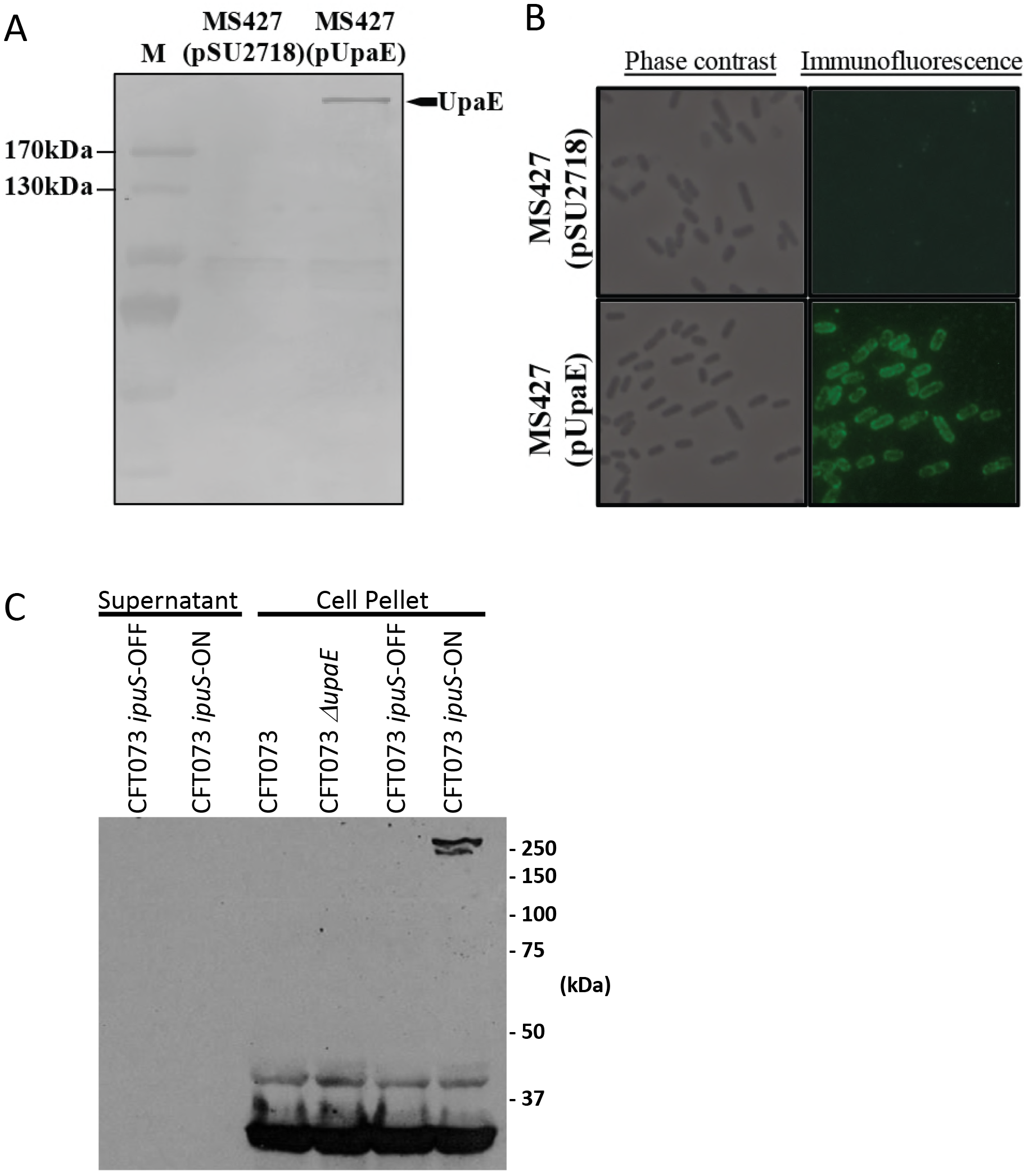
Expression and surface localisation of UpaE in *E. coli* K-12. (A) Western blot analysis of whole cell lysates prepared from MS427(pSU2718) vector control and MS427(pUpaE). A band corresponding to UpaE (271 kDa) was detected in MS427(pUpaE) but not in the MS427(pSU2718) control. Lane M refers top molecular weight markers; the 170 kDa and 130 kDa proteins are indicated. (B) Phase contrast and immunofluorescence microscopy using specific antisera against UpaE. Positive reactions indicating the surface localisation of UpaE were detected in MS427(UpaE) (bottom), but not in the MS427(pSU2718) vector control (top). (C) Western blot of pelleted cells solubilized in crack buffer or concentrated 10ml TCA precipitations of culture supernatants. Protein detection was performed using a polyclonal anti-sera raised to a UpaE-MBP fusion protein. A band consistent with the predicted 271 kDa size of UpaE is only present in the cellular fraction of phase lock on cells.

### UpaE mediates biofilm formation and adhesion to ECM proteins

After assessing expression and localization, we probed the functionality of UpaE. As surface-bound autotransporters are frequently involved in biofilm formation or adherence, we assessed biofilm production. The parent strain MS427 is unable to form biofilms, but introduction of the plasmid born copy of UpaE resulted in a significant increase in biofilm production when assessed by crystal violet staining (Fig. 6A). We also investigated the ability of UpaE to mediate adherence to human ECM proteins. Adherence to MaxGel, a commercially available mixture of collagens, laminin, fibronectin, tenascin, elastin and a number of proteoglycans and glycosaminoglycans, was significantly increased in the UpaE overexpression strain compared to empty vector controls (Fig. 6B). Further examination revealed UpaE mediates significant adherence to fibronectin, laminin, and collagen I, II, V specifically (Fig. 6B). Together these results suggest that UpaE is a surface exposed autotransporter and facilitates both biofilm formation and adherence to human ECM proteins.

**FIG 6.**
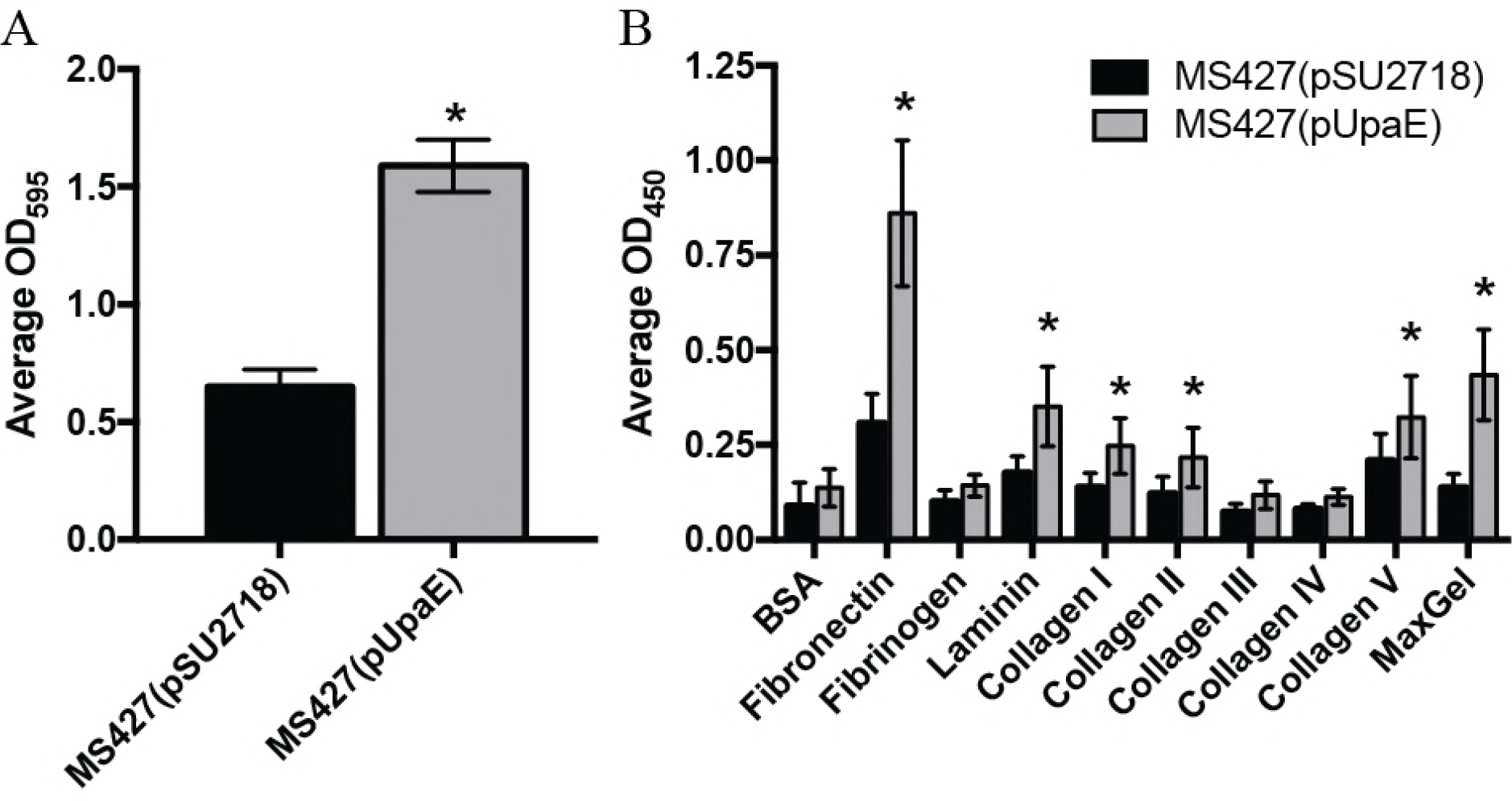
UpaE mediates biofilm formation and adhesion to ECM proteins. (A) Polystyrene 96 well microtiter plates (Corning) were used to monitor biofilm formation. Overnight cultures were subcultured 1/100 into fresh M9 minimal media supplemented with 1mM IPTG, and incubated shaking at 37oC for 24h. Biofilms were stained with a 0.1% crystal violet solution, and quantified by dissolving the crystal violet with an acetone-ethanol mix and measuring the OD595. MS427(pUpaE) was able to form a better biofilm as compared to the vector control. Experiments were performed in triplicate. (*; P < 0.05, unpaired student t-test). (B) MS427(pSU2718) vector control and MS427(pUpaE) were incubated in microtiter plates coated with ECM proteins. Non-adherent bacteria were washed off and remaining bound cells were detected with a specific E. coli antiserum. MS427 overexpressing UpaE bound to fibronectin, laminin, collagen I, II and V, and MaxGel. Experiments were performed in triplicate. (*; P < 0.05, unpaired student t-test).

### Phenotypes of the *ipuS* ON vs OFF phase states in CFT073

Due to the conservation of this region in many UPEC strains and the known links between adherence promoting autotransporters and phase variation in pathogenesis (20, 21, 54–58), we predicted that inversion would play a role in virulence related phenotypes. We previously demonstrated that in the *fimS* ON state that there is a reduction in motility compared to the OFF position (23). Overnight liquid cultures were used to inoculate the surface of Adler’s Motility Medium agar plates and diameters of the swimming zones were measured after ~21hrs of growth at room temperature. We observed that the *ipuS* OFF state is more motile than the *ipuS* ON state in a Type 1 pili OFF background (Fig. 7A). The same trend was observed in the Type 1 pili ON background, however the non-motile nature of Type 1 pili ON cells made it difficult to clearly discern the *ipuS* effects (Fig. 7A).

**FIG 7.**
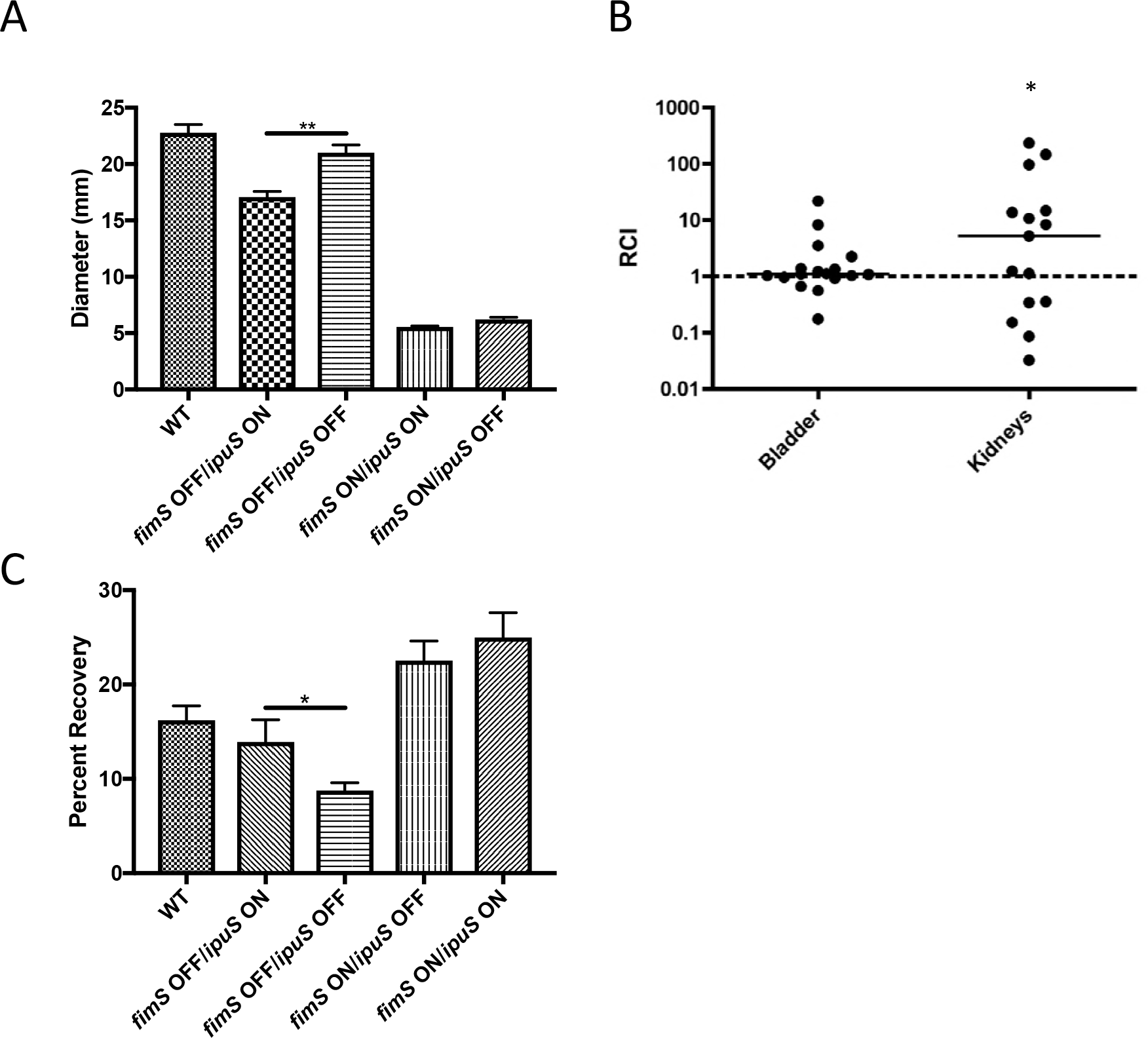
Phenotypic readouts of the *ipuS* phase state in a CFT073 background. (A) Swimming motility of phase locked strains. 1μl of liquid culture normalized to OD600 = 0.01 of each of the strains WAM2266 (WT), WAM5088 (*ipuS ON/fimS* OFF), WAM5063 (*ipuS OFF/fimS* OFF), WAM 5064 (*ipuS* ON, *fimS* ON), and WAM5065 (*ipuS OFF/fimS* ON) was inoculated in the center of 7 plates containing Adler’s motility medium. Values represent diameter of the swimming zone after ~21 hr of room temperature incubation (Mann-Whitney, p<0.005). (B) 72HR competitive infection of WAM5064 (*fimS* ON, *ipuS* ON) vs. WAM5146 (*fimS* On, *ipuS* OFF, *lacZYA*). Relative competitive indices were calculated from bladder and kidney homogenates at 72hpi with lines representing the medians. WAM5064 (*ipuS* ON) has a 5-fold advantage in the kidneys (*p<0.05, Wilcoxon Signed Rank Test) at 72hpi. (C) Adherence to A-498 human kidney epithelial cells. WAM2266 (WT), WAM5088 (*ipuS ON/fimS* OFF), WAM5063 (*ipuS OFF/fimS* OFF), WAM 5064 (*ipuS* ON, *fimS* ON), and WAM5065 (*ipuS OFF/fimS* ON) were allowed to adhere to monolayers of A-498 cells then unbound bacteria were removed by washing. In a *fimS* OFF background, turning *ipuS* ON causes a 5% increase in adherence (WAM5088 vs. WAM 5063) (p<0.05, t-test).

### *ipuS* orientation affects colonization of the kidneys in a murine model of UTI and adherence to human kidney epithelial cells

We next assessed if the *ipuS* phase state results in a difference in colonization in the murine model of UTI. To test this, we performed competition assays using the *ipuS* locked ON and OFF strains to address putative UpaE dependent effects on colonization. A Δ*lacZYA* mutant variant of the *ipuS* locked OFF strain (WAM5146) was used to facilitate generation of competitive indices using MacConkey lactose medium. Previous experiments indicated that a Δ*lacZYA* mutant of CFT073 competes equally against WT CFT073 (59). 50μl inoculums containing an equal ratio of the *ipuS* locked ON and OFF strains (totaling 10^8^ CFU) were transurethrally delivered into the bladder of 6-week-old female CBA/J mice and the infections were allowed to progress for 72 hours. The animals were sacrificed and their bladders and kidneys excised, homogenized, and plated on MacConkey’s lactose medium. Ratios of ON/OFF bacteria at sacrifice were normalized to the input ratio to generate relative competitive indices (RCI). The Type 1 fimbriae locked ON variants of the *ipuS* ON/OFF strains were used in the experiment because Type 1 fimbriae deletion strains are severely attenuated in mouse models of UTI (15). At 72hpi, a 5-fold advantage (p<0.05) for the *ipuS* ON state was observed in the kidneys (Fig. 7B). No difference was seen in the bladder at 72hpi (Fig. 7B). To further assess the role of UpaE in facilitating infection *in vivo*, we also performed competition assays between WT CFT073 and an otherwise isogenic *ipuR/upaE* mutant. However, in this context no significant competitive difference was observed in the bladder or kidneys (Fig S1). We postulate this is due to the phase permissive background of the *ipuR/upaE* mutant strain. Locking *fimS* ON to help facilitate competitive infections in the ipuS ON/OFF strains also suppresses the production of P pili (41), a kidney specific adhesion factor. We predict that the ability of the *ipuR/upaE* mutant to produce P pili compensates for the difference observed in the *ipuS* OFF vs. ON strains.

The locked strains that were used in the competitive infection assays were virtually non-motile due to the constitutive expression of Type 1 pili (23). Importantly, this suggested that the in vitro swimming motility defect of WAM5088 was not the cause of the *ipuS* dependent kidney colonization advantage. Rather, we speculated that the change in colonization was at least partially attributed to UpaE expression and its effect on adherence. To determine if the adhesive properties of UpaE may have contributed to the kidney specific advantage observed in vivo, we assayed the four *ipuS/fimS* phase locked strains for their ability to adhere to human kidney epithelial cells. The strains were incubated with confluent monolayers of A-498 cells (MOI=10) for 1hr and adherence was assessed by direct determination of colony forming units. The number of adherent bacteria were normalized to the input and expressed as percent adherence. Locking *ipuS* ON in a *fimS* OFF background increased adherence to kidney epithelial cells (p<0.05) suggesting that expression of UpaE facilitates adhesion to the kidney epithelium. Type 1 pili also promoted kidney adherence, however locking both switches on did not cause a synergistic increase in adhesion (Fig. 7C).

## Discussion

Phase variation is defined as rapid and reversible ON/OFF changes in gene expression (60). It occurs by several different molecular mechanisms and contributes to virulence in multiple pathogens including *E. coli*, *Neisseria meningiditis*, *Mycoplasma agalactiae, Listeria monocytogenes*, and *Clostridium difficile* (61–66), It is an advantageous form of gene regulation for pathogens as it helps a population cope with sudden changes in environmental conditions during infection (67). The presence of a subset of the population in alternative phase states circumvents the need for transcriptional and translational activation steps in response to changing conditions. CFT073 has two known phase variable elements, *fimS* and *hyxS* (23, 45). Here, we identified a third phase variable element (*ipuS*). We demonstrate that the orientation of the *ipuS* element in CFT073 controls the transcription of two downstream genes (*ipuR* and *upaE*), which in turn affects motility and kidney colonization in mice. Additional analysis of UpaE revealed it is surface localized and mediates biofilm formation and adhesion to ECM proteins.

When comparing the sequences of the half sites, *ipuS* appears to be an intermediate between *fimS* and *hyxS* (Fig. 2). This suggests that the non-proximal recombinases FimB, FimE and FimX would have activity at *ipuS.* Indeed, we found that FimE and FimX are catalytically active at *ipuS* (Fig. 3). Only limited sequence similarity is present between *fimS* and *hyxS* in UTI89 (45), which may account for why only the proximally encoded FimX is able to function at *hyxS* in both CFT073 and UTI89.

Though the assay described in this work was not explicitly quantitative, the five recombinases display apparent differences in their efficiency for inversion of the *ipuS* element, which indicates potential directional biases (Fig. 3). A directional bias for FimB/FimE at *fimS* has been characterized extensively in *E. coli* K-12 and is due to sequence specificity of the recombinases at the inverted repeats and surrounding sequence (22, 68–71). FimE is unable to bind to the *fimS* half sites in the OFF orientation, which restricts its activity for catalyzing ON to OFF inversion. By mutating the regions outside the inverted repeats to resemble the ON or OFF state, this specificity can be reversed (69). It is possible that the apparent decreased efficiency of FimE and IpuB at *ipuS* is due to a defect in their ability to bind to the template. Electrophoretic mobility shift assays have been performed with FimB/FimE at *fimS* to characterize this effect, however the recombinases are notoriously difficult to purify, complicating the analyses (70, 71). Further studies focused on precise assessment of catalysis and inversion frequencies, such as the application of read-mapping approaches based on deep sequencing to monitor switching (46), are needed to assess how phase bias at *ipuS* may contribute to population polarization.

The orientation of *ipuS* may also directly influence *fimS* or *hyxS* orientation, but such effects were masked by the need to lock all three switches in our analysis. Other investigators have generated *fimS* locked strains by mutating the sequence of the inverted repeats (64). Using this approach would facilitate locking *ipuS* orientation while permitting inversion of the other two elements, helping to identify *ipuS* effects at *fimS* and *hyxS*. However, the five recombinases recognize the inverted repeats in a sequence specific manner (69–71) so manipulating the local sequence may inherently change recombinase-binding affinity. In the context of a complete network, where multiple sites compete for limited quantities of each recombinase, changing the half sites could perturb the orientation of the other switches by altering recombinase availability. As such, it stands to reason that the orientation of all three elements is interrelated, as they compete for a limited pool of shared enzymatic machinery.

5’RACE analysis indicated the presence of a transcriptional start site in the *ipuS* element (Fig. 4). The promoter is part of the full-length *ipuA* coding region and reorientation of the element turns transcription of *ipuR/upaE* ON/OFF. By sequence inspection for conserved promoter motifs proximal to the transcription start site we were able to identify a putative *rpoD* dependent promoter. Direct in vitro transcription assays using RNA polymerase holenzyme are planned for the future to support this supposition.

*ipuR* is a predicted LuxR-type transcriptional regulator. LuxR-type regulators, are two domain proteins that contain an autoinducer and DNA binding domain. They have been implicated in virulence of multiple pathogens, including *Vibrio* spp., several classes of pathogenic *E. coli*, and *Mycobacterium tuberculosis*, where they often regulate systems involved in biofilm formation and motility (45, 55, 56, 72). The regulon sizes of these proteins are highly variable. Some regulate one or a few specific targets while others have much broader effects (55). The effects of *ipuS* described here in murine infection models, tissue culture, and in vitro systems appear UpaE dependent. It remains unclear what role IpuR plays, if any, in the regulation of *upaE* or other target genes. While we did not observe ipuR-dependency in the phenotypes described here we also cannot rule out contribution to these or other putative phenotypes. Definition of the *ipuR* regulon and its contribution to UPEC biology and pathogenesis are active areas of research.

Autotransporters are large multi-domain proteins that belong to the Type V secretion system (73). They possess an N-terminal signal sequence that targets the protein to the Sec machinery for transport into the periplasm, a passenger domain that is either secreted or cell-surface associated, and a C-terminal translocator domain that is embedded in the outer membrane and helps facilitate translocation of the passenger domain (74–76). CFT073 possesses genes encoding multiple different autotransporters, which function either as adhesins or secreted toxins (20, 21, 51, 57, 77, 78). One well studied autotransporter is Ag43, a surface bound protein found in most *E. coli* strains, is phase variable mediates cell-cell adhesion, biofilm formation and long-term colonization of the mouse bladder (49, 77, 79). Ag43 phase variation is mediated by the combined action of DAM methylase (activation) and OxyR (repression) (80–82). Additionally, altered methylation patterns in key regions modulate Ag43 transcription and expression of Ag43 is important for facilitating infection in the murine model (77). Here we characterize a previously uncharacterized autotransporter, UpaE, which represents another phase variable autotransporter of *E. coli*. We show that UpaE is surface exposed, mediates biofilm formation and adherence to human ECM proteins. Our data also imply that UpaE enhances UPEC virulence based on analysis of an *ipuS* locked ON strain in mice. Importantly, we previously observed that the *ipuA-upaE* region is more prevalent in UPEC (37%) than commensal strains (7%) suggesting this system to be a relevant virulence mechanism for many UPEC strains (23). Further studies confirming the adhesive properties of UpaE and the conditions/factors that select for its expression are in progress.

To assess the role of *ipuS* in virulence in the murine model of UTI we infected female mice transurethrally into the bladder. We assessed colonization of the bladder and kidneys in a mixed competitive infection assay using *ipuS* locked ON and OFF strains, and in a type 1 fimbriae locked ON background (Fig. 7B). Locking type 1 fimbriae ON helps facilitate consistent infections as lock OFF strains are severely attenuated (15). However lock ON strains have impaired swimming motility, which is also important for colonization, and type 1 fimbriae expression inhibits the production of other adhesive pili including the kidney specific P pili (41, 83). The interrelated nature of these systems makes it difficult to study their effects in isolation and may also account for the high degree of variability observed in animal models. Further development of phase locked *ipuS* strains that are decoupled from *fimS* and *hyxS* inversion are underway to evaluate *ipuS* specific effects.

Tyrosine recombinases often function at invertible elements encoded in close proximity to themselves (44). However, there are exceptions to this generalization. For example, in depth analysis of *Bacteriodes fragilis* has revealed extensive networks of switches and recombinases that function at local and distant sites in the chromosome (84–88). One such enzyme, Mpi, can catalyze inversion of 13 elements located throughout the *B. fragilis* chromosome (88). This inversion network controls the expression of surface architecture components and is predicted to function as a mechanism for global surface remodeling in response to changing environmental conditions (85, 88). The identification of *ipuS*, demonstrated recombinase cross reactivity among the three invertible elements, and known environmental stimuli that influence inversion of the switches (38, 39, 89, 90) suggests the existence of a complex network in UPEC (23, 45). UPEC encounter a variety of different conditions during colonization of a human host, for example in the gut, urethra, bladder, kidneys and bloodstream. We hypothesize that population heterogeneity generated by multiple mechanisms, including differential gene regulation, epigenetic regulation, and the phase variable network described here provide a means for UPEC to successfully colonize these different environments.

## Methods and Materials

### Bacterial strains, cell lines, plasmids and culture conditions

All of the strains, cell lines and plasmids used in this study are listed in Table 1. In frame deletion mutants of CFT073 were generated using a modification of the Lambda-Red method of homologous recombination to include phage transduction of the marker into a clean genetic background by ϕEB49 prior to removal of the cassette via pCP20 (91, 92). Phase lock mutants were generated by sequential deletion of the five previously described tyrosine recombinases in CFT073 (23). Upon deletion of the final recombinase, multiple colonies were screened to identify mutants with all four possible combinations of *fimS* and *ipuS* phase states. *lacZ* transcriptional fusions were generated using methods described previously with the suicide vector pFUSE (93). All strains were cultivated in Luria Bertani (LB) broth, LB agar, or on MacConkey lactose medium unless otherwise indicated. Antibiotic selection employing kanamycin (50μg/ml), chloramphenicol (20μg/ml), or carbenicillin (250μg/ml) was used as appropriate.

The kidney epithelial cell line A-498 (ATCC HTB-44) was grown in RPMI 1640 with L-glutamine (Mediatech, Inc.) supplemented with 20% fetal bovine serum (Atlanta Biologicals, Lawrenceville, GA), 10mM HEPES, and 1mM sodium pyruvate (Mediatech, Inc.). Cells were grown at 37°C with 5% CO_2_ and utilized at less than 10 passages.

### *ipuS* switch state analysis

The *ipuS* region was amplified by PCR using GoTaq Green Master mix (Promega) from 0.5μl of overnight LB broth cultures using the forward primer 5’-GTGGCGATGGGAAGGAAACG-3’ and reverse primer 5’-AAAACCCCGCCAACGCATACTC-3’. Thermocycling conditions were 94°C for 2min, 25 cycles of 94°C for 30sec, 57°C for 30sec, 72°C for 1min 30sec, and 72°C for 7min. The resulting 1289 bp product was purified using a Qiaquick PCR purification kit (Qiagen) and digested with PacI (New England Biolabs). Digested fragments were electrophoresed through 2% agarose gel and stained with ethidum bromide. Sizes of the restriction products correspond to the state of the switch (407 bp and 882 bp ‒ phase ON; 186 bp and 1103 bp, phase OFF).

### Construction of plasmids

The *fimB*, *fimE*, *fimX*, *ipuA* and *ipuB* CFT073 recombinases were cloned into either pACYC177 or pACYC184. For constructs built within the pACYC177 backbone, the respective recombinase genes were constitutively expressed from the plasmid encoded kanamycin resistance gene promoter. For constructs built within the pACYC184 backbone, the respective recombinase genes were expressed from their native promoter. The *upaE* gene was amplified from CFT073 with primers 7799 (5’- GACCTGCAGGCATGCAAGCTATGAAGGAGGAGTGGTATGAATAAAGTATATAAAG -3’) and 7800 (5’- CGACGGCCAGTGCCAAGCTTTAGAATATATATTTAATACC -3’), and inserted into pSU2718 using a modified ligation-independent cloning protocol (94). Briefly, the pSU2718 plasmid was digested with HindIII, and both cut plasmid and PCR product were treated with T4 polymerase to generate complementary overhangs. The T4 polymerase-treated insert and plasmid were mixed in a 3:1 ratio, and incubated on ice for 30 minutes to generate pUpaE. All plasmids were confirmed by PCR and sequencing of the inserts.

### 5’RACE of *ipuS* element

The *ipuS* transcription start site was identified using 5’ Rapid Amplification of cDNA Ends (5’RACE) (Invitrogen). Gene specific nested primers were designed according the manufacturer’s instructions. RNA was extracted from 1ml of log phase (OD_600_=0.5) culture of WAM5064 and WAM5065 using Trizol reagent (Invitrogen). Contaminating DNA was removed by on column DNase treatment and Pure Link RNA spin column purification (Invitrogen, Grand Island, NY) and the resulting purified RNA samples were stored in nuclease-free water at −20°C. Aliquots of the isolated RNA were processed using the 5’RACE Kit and gene specific primers (Invitrogen) according to manufacturer’s instructions. The resulting PCR products were sequenced using Sanger dideoxy chain termination sequencing to identify putative transcription start sites.

### UpaE polyclonal antibody production and western blotting

Rabbit polyclonal anti-UpaE serum was raised to a recombinant maltose binding protein *malE-upaE* gene fusion using the pMal-p2x vector (New England Biolabs). Residues S24 to G2000 of UpaE were present in the fusion protein. Expression of the fusion protein was induced by addition of IPTG to the growth medium. Inclusion bodies containing the large fusion protein were solubilized in crack buffer (2% sodium dodecyl sulfate [SDS]–10% glycerol–5% p-mercaptoethanol‒1 mM bromophenol blue‒62 mM Tris)) subjected to SDS-polyacylamide gel electrophoresis (SDS-PAGE). The large Coomassie-stained fusion protein band was excised from the gels and then used as an immunogen in rabbits.

### In vivo Expression of UpaE was determined by western blot

Cell pellets were solubilized in crack buffer and subjected to SDS-PAGE in 10% gels. Concentrated culture supernatants were prepared by taking 10 mls of filtered late log-phase L-broth adding tricholroacetic acid to make a final 10% concentration. After overnight incubation at 0-4 °F, precipitates were collected by centrifugation and solubilized in 20 μls of crack buffer. 1M Tris in 1 μl volumes were added until the resuspended pellet changed from yellow to blue in color. Protein detection was performed using the primary UpaE polyclonal antibody described above, secondary anti-rabbit-HRP (Bio-Rad), and chemiluminescent detection by Amersham ECL Prime Western Blotting Kit (GE Healthcare).

### Immunofluoresence microscopy

Immunofluoresence microscopy was performed essentially as previously described (53). Overnight cultures supplemented with the appropriate antibiotics and 1mM of IPTG were fixed to an OD_600_ of 0.4, spotted onto a glass slide and allowed to dry. The cells were fixed with 4% paraformaldehyde (PFA), washed with PBS, and blocked with 0.5% BSA. The slides were incubated with the anti-UpaE antibody, washed with PBS, and further incubated with a secondary goat anti-rabbit antiserum couple to fluorescein isothiocyanate (FITC). The slides were washed, air-dried and mounted with ProLong Gold (Invitrogen), and examined under a ZESS Aixoplan 2 epifluorescence microscope.

### Biofilm Assay

PVC 96-well microtiter plates (Corning) were used to monitor biofilm formation as previously described (95). Briefly, cells were grown for 18h in LB at 37°C, washed to remove unbound cells, and stained with 0.1% crystal violet. Quantification of the cells was performed by dissolving the crystal violet with ethanol-acetone (80:20) and taking the absorbance reads at OD_595_. Results were presented as the mean of eight replicate wells from three independent experiments. The data were analyzed using the unpaired student t-test with Graphpad Prism 7 software. The graph represents results of three independent experiments with standard deviations included.

### ECM adhesion assay

Bacterial binding to ECM proteins was performed in a microtiter plate enzyme-linked immunosorbent assay (51). Briefly, microtiter plates (Maxisorp; Nunc) were coated overnight with MaxGel human ECM (10μg/ml) or 2μg/ml of collagen (types I-V), fibronectin, fibrinogen, laminin or bovine serum albumin (BSA)(Sigma-Aldrich). Wells were washed with TBS (137 mM NaCl, 10 mM Tris, pH7.4) and blocked with TBS-2% milk for 1 h. Bacterial cultures were standardized to OD_600_ of 0.1 in TBS, and 200μl of the cultures were added to the plates. After washing to remove unbound cells, adherent bacteria were fixed with 4% PFA, washed, and incubated with an anti-*E. coli* serum (Meridian Life Sciences Inc.) for 1 h. The cells were washed, and incubated with a secondary anti-rabbit-conjugated horseradish peroxidase antibody for another 1 h. Following a final wash, adherent bacteria were detected by adding 50μl of TMB. After 15 min, 50μl of 1M HCl was added to stop the reaction, and the absorbance was read at OD_450_. The data were analyzed using the unpaired student t-test with Graphpad Prism 7 software. The graph represents results of three independent experiments with standard deviations included.

### Swimming motility assay

Strains were grown in LB broth with appropriate selection over night at 37°C with shaking. The overnight cultures were normalized to OD_600_= 0.5 and 1μl was inoculated directly into the center of a Petri plate containing 20ml of Adler’s Motility Medium (0.3% agar, 0.5% NaCl, 1.0% tryptone) (N=7). Plates were incubated lid side up at room temperature for approx. 21 hours. The diameter of the zone of swimming was measured twice at perpendicular angles for each plate and the averages were plotted. The data were analyzed using the data analysis software package Prism (Graphpad) to determine statistically significant differences (p<0,05) between strains by Mann-Whitney test.

### Kidney epithelial cell adherence assay

A-498 cells were seeded into 12-well plates at 2.5 × 10^5^ cells/well and grown to near confluence. Monolayers were washed two times with assay medium (serum and antibiotic free culture medium) and preincubated for 20min at 4°C in 1ml assay medium. Triplicate wells were inoculated with bacteria (MOI=10) and were settled onto host cells by centrifugation at 500 = g for 5min. After 1hr incubation at 4°C, monolayers were washed three times with HBSS (Hyclone), incubated for 5min at 37°C in 500μl 0.025% trypsin/0.03% EDTA in HBSS, lysed with 0.1% Triton X-100 in ddH_2_O and plated on LB-agar plates. Adherence was calculated as the ratio of the number of bacteria recovered to the number of bacteria inoculated into each well and expressed as percent adherence The data were analyzed using the data analysis software package Prism (Graphpad) to determine statistically significant differences between strains by unpaired students t-test.

### Murine model of UTI

Six-week-old female CBA/J (Harlan Laboratories,) mice were used for all infections. Cells were grown in static LB-broth and infections were performed as described previously (59, 96). For competitive infections WAM5146, a *lacZYA* mutant variant of WAM5065, was used to facilitate generation of competitive indices with MacConkey’s lactose medium. Previous experiments indicate that *lacZYA* activity has no influence on colonization in the murine model of UTI (97). When using locked strains to examine the effects of *ipuS* variable phase states in the mouse, the type 1 fimbriae lock ON versions were used in order to facilitate consistent infections as type 1 deficient UPEC strains are severely attenuated (15). Bars on presented data indicate the medians of the non-Gaussian distributed data sets. Wilcoxin-Signed Rank Tests were performed with Prism (Graphpad Software, La Jolla, CA) to determine statistical significance and reported when p<0.05. This study was done in strict agreement with the recommendations found in the Guide for the Care and Use of Laboratory Animals of the National Institutes of Health. The murine model UTI protocol was approved by the UW-Madison Animal Care and Use Committee (Permit Number: M00450-0-07-08).

## Acknowledgements

This work was supported by National Institutes of Health (NIH) grant R01-DK063250-07, grants from the National Health and Medical Research Council (NHMRC) of Australia, a Robert Turell Professorship awarded to R.A. Welch, and an NHMRC Senior Research Fellowship awarded to M.A. Schembri.

---

**FIG S1.**
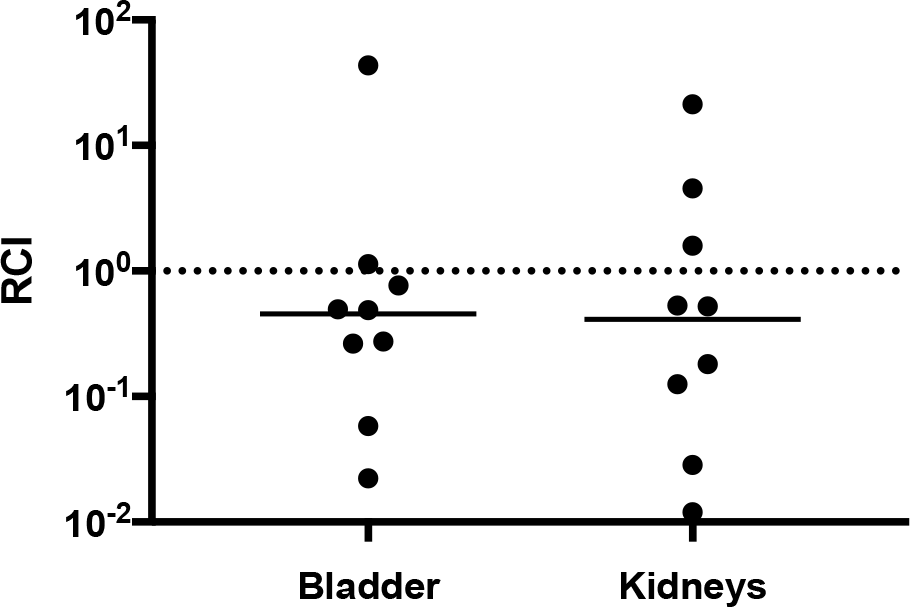
72HR competitive infection of WAM5048 (CFT073 *ΔipuR/upaE*) vs. WAM4520 (CFT073 *ΔlacZYA*). Relative competitive indices were calculated from bladder and kidney homogenates at 72hpi with lines representing the medians.

